# Dynamic Electrode-to-Image (DETI) Mapping Reveals the Human Brain’s Spatiotemporal Code of Visual Information

**DOI:** 10.1101/2021.02.19.431983

**Authors:** Bruce C. Hansen, Michelle R. Greene, David J. Field

## Abstract

A chief goal of systems neuroscience is to understand how the brain encodes information in our visual environments. Understanding that neural code is crucial to explaining how visual content is transformed via subsequent semantic representations to enable intelligent behavior. Although the visual code is not static, this reality is often obscured in voxel-wise encoding models of BOLD signals due to fMRI’s poor temporal resolution. We leveraged the high temporal resolution of EEG to develop an encoding technique based in state-space theory. This approach maps neural signals to each pixel within a given image and reveals location-specific transformations of the visual code, providing a spatiotemporal signature for the image at each electrode. This technique offers a spatiotemporal visualization of the evolution of the neural code of visual information thought impossible to obtain from EEG and promises to provide insight into how visual meaning is developed through dynamic feedforward and recurrent processes.

## Introduction

Upon viewing a new scene, the brain transforms the ambient light array hitting the retinae into semantically meaningful content that enables intelligent behavior, all within the first 300 ms of viewing. However, the ensuing series of representational transformations that shape our understanding of the visual world are not well-understood. At the most fundamental level, we know that visual information is processed differentially by multiple neural populations that act like nonlinear filters, each coding for specific types of information in narrow bands of spatial frequency and orientation [1–3]. Further, because real-world environments are broadband in both spatial frequency and orientation, each location within a given scene will simultaneously activate a range of tuned visual neurons [4–10]. Given this, recent efforts have used visual filter-based encoder models to predict fMRI-defined patterns of blood oxygen-level dependent (BOLD) activity based on real-world scene inputs [11–13]. Such voxel-wise encoder models of BOLD signals have provided insight into the nature of how humans internalize external information, and on how the neural code in early visual cortex maps onto and supports higher level semantic representations [14–17]. However, such insight is based on a static view of the visual code (due to the temporal limitations of fMRI). The neural code for visual information is highly dynamic as local and long-range recurrent processes act to change the local activity patterns that are evoked by scenes over a short period of time [18–20]. However, these time-varying transformations are not yet well-characterized, thus hindering our understanding of how they enable the construction of a meaningful representation of our visual world.

Traditional electroencephalography (EEG) has excellent temporal resolution and has been used as an attempt to characterize the time-varying nature of visual information [21–24]. Nonetheless, EEG suffers from scalp interference and dipole cancellation on the scalp, so these efforts have only provided a very coarse estimate of time-varying scene transformations, with virtually no insight into how local scene information is encoded and transformed over time. This is unfortunate because the early spatiotemporal transformations of visual information likely serve to help shape how our visual world is represented in many higher-level cortical networks, ultimately shaping scene-related semantic processes [14,21, 25–28]. For instance, multiple networks across lateral occipitotemporal, dorsal, ventral temporal, and medial temporal cortices have all been shown to possess a retinotopic organization and relative selectivity to different spatial frequencies [29–35].

In this study, we introduce dynamic electrode-to-image (DETI) mapping: an analytical approach to map time-varying neural signals from visual evoked potentials (VEPs) to every pixel location of complex real-world scenes. This technique therefore offers the ability to visualize and analyze the spatiotemporal evolution of visual encoding across the early stages of visual processing. To circumvent the problems inherent in EEG measures, we mapped the localized outputs of a spatial frequency tuned log-Gabor encoding model to different VEPs within a geometric state-space framework. Specifically, we measured the correspondence between the high-dimensional output variation produced by our encoding model at every location within large-field visual scenes and the response variation of VEPs measured at each electrode across the posterior region of the scalp. At its heart, our method reduces the dimensionality of VEP signals measured at each electrode at different points in time and then maps those signals via an encoding model to each pixel within and across a relatively large set of images. This geometric state-space mapping procedure enables the mapping to take place across large sets of scenes (**Figure 1**) as well as for individual scenes (**Figure 2**). Specifically, this method provides 1) a general spatiotemporal view of scene encoding over an entire set of images, thereby allowing a visualization of the general coding strategy over time, and 2) a scene-specific spatiotemporal view to visualize the various transformations that each scene undergoes over time. Further, this technique offers a rich source of spatiotemporal data to explore a wide variety of questions concerning the various transformational states of visual coding once thought impossible to address with EEG measures.

**Figure 1.**
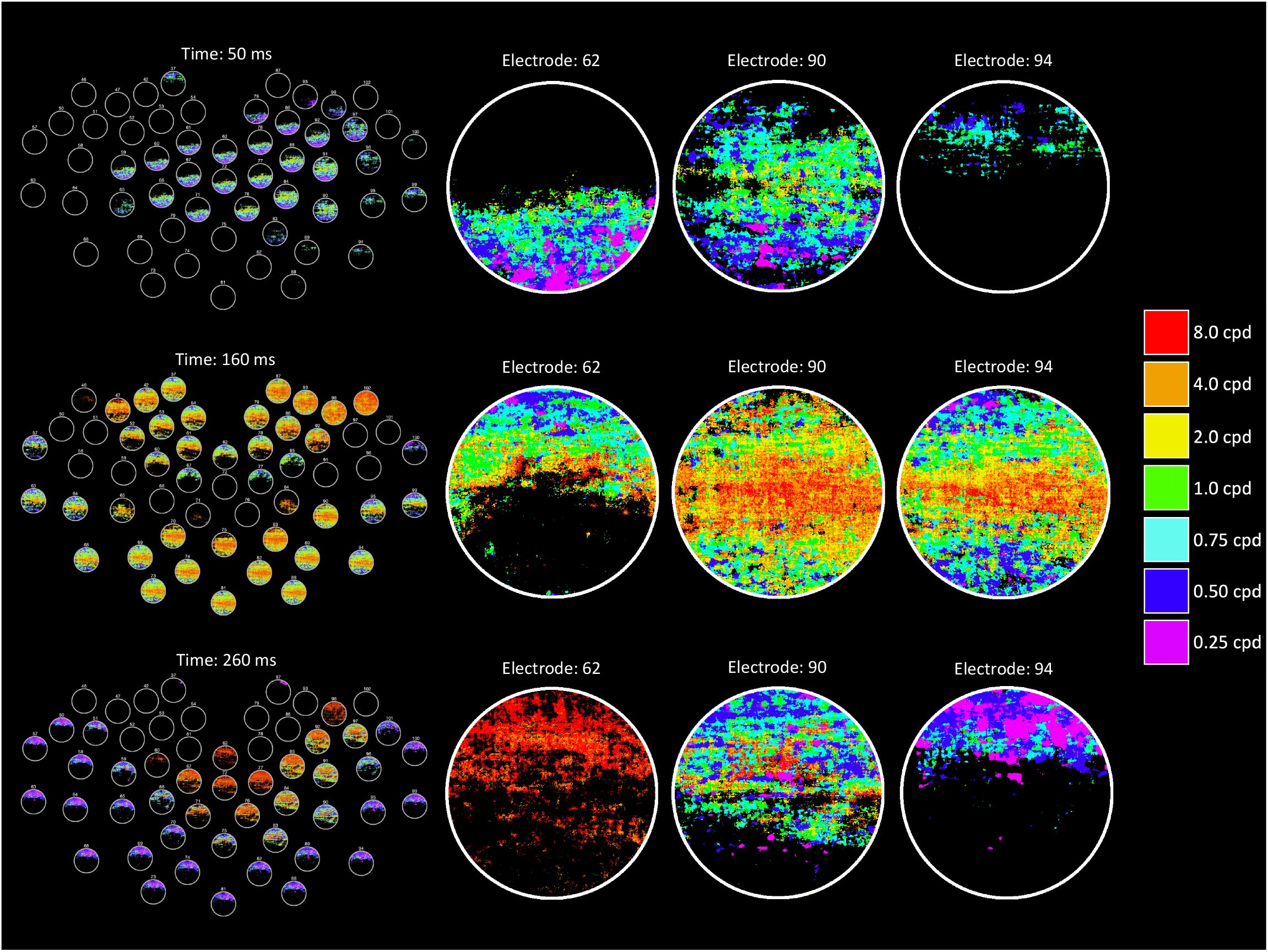
Example DETI maps from the image-general analysis at different time points. The movie version of this figure can be downloaded here [<u>**LINK**</u>]. The left-hand column shows a topographical map of the posterior electrodes, illustrating the variation of DETI maps across that scalp region. On the right-hand side, each column shows the spatiotemporal evolution of the visual code for different electrodes (each row corresponds to the time given on the left-hand side). The colorbar shows the spatial frequency tuning peak (in cycles per degree; cpd) of the encoder that was mapped to each pixel in the DETI maps. Note that the maps are circular because the stimuli were windowed with a circular window (see Materials & Methods).

**Figure 2.**
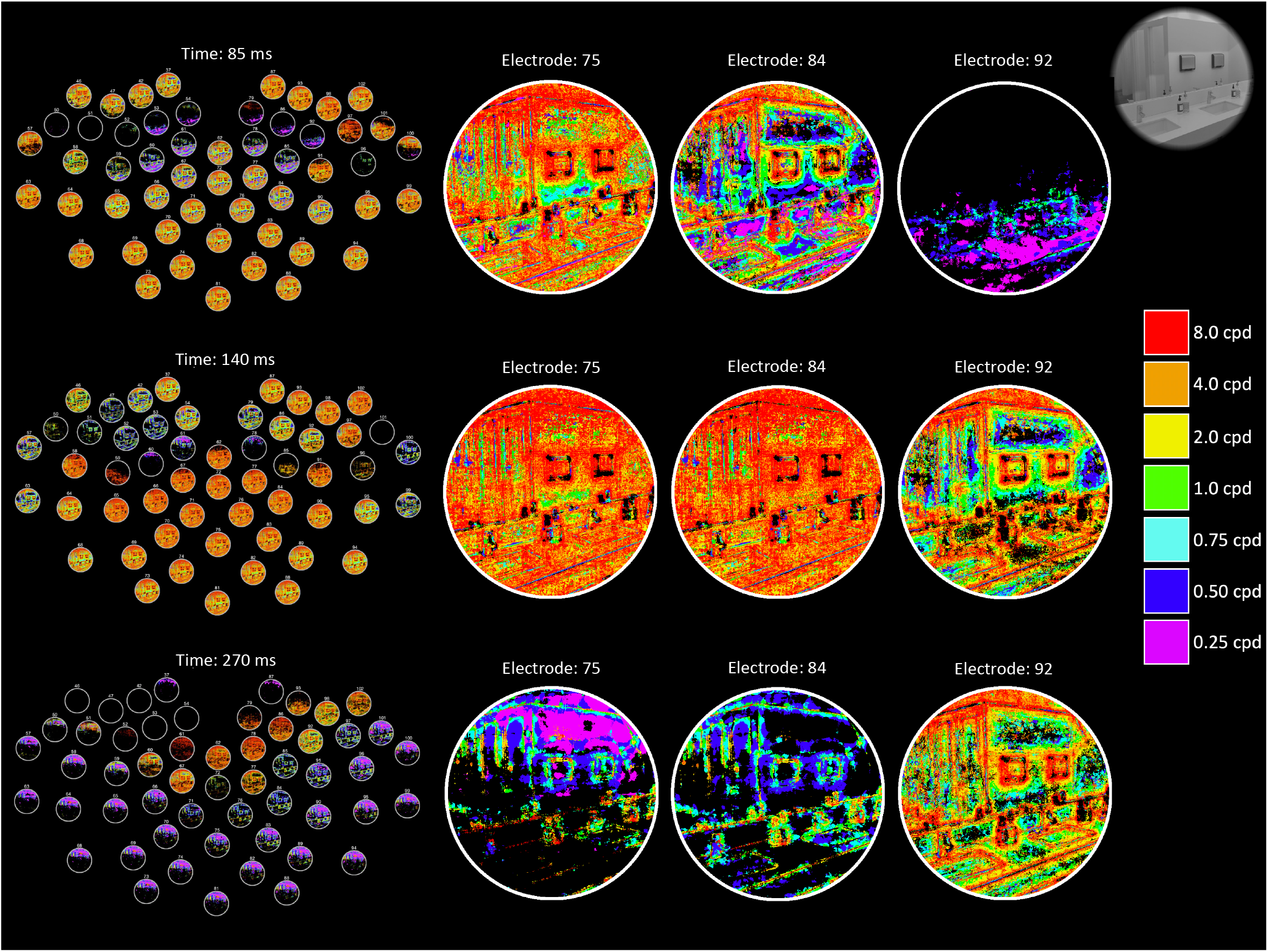
Example DETI maps from the image-specific analysis at different time points. The movie version of this figure can be downloaded here [<u>**LINK**</u>]. The layout and details of this figure are identical to **Figure 1**, except here we are showing the spatiotemporal evolution of the visual code for an example image (shown in the upper right-hand corner).

In this report, we took advantage of the local coding abilities of this approach to characterize the transformational states of scenes. That characterization revealed that scenes undergo a series of nonuniform transformations that prioritize different spatial frequencies at different regions of scenes. Further, the spatiotemporal visual code varies in a location specific manner, likely reflecting the underlying principles of the early visual code, thereby offering some perspective on how each transformational state informs and refines the conceptual meaning of our visual world.

## Results

The DETI mapping procedure can be broken down into three pipeline operations followed by two different mapping procedures (illustrated in **Figure 3**). The first pipeline operation projects high-dimensional VEP data into a lower dimensional space via time-resolved principal component analysis (PCA). The second projects each stimulus image to each of seven log-Gabor encoders on a pixel-by-pixel basis. The third and final pipeline operation links the lower-dimensional VEP data to each pixel in the encoder space. From there, DETI maps can be constructed for each electrode based on all scene stimuli (an image-general analysis) or for each scene stimulus (image-specific analysis). The following subsections outline each DETI pipeline operation, followed by the results of the DETI mapping procedure at two levels of analysis. The code for both mapping operations can be downloaded here [<u>**LINK**</u>].

**Figure 3.**
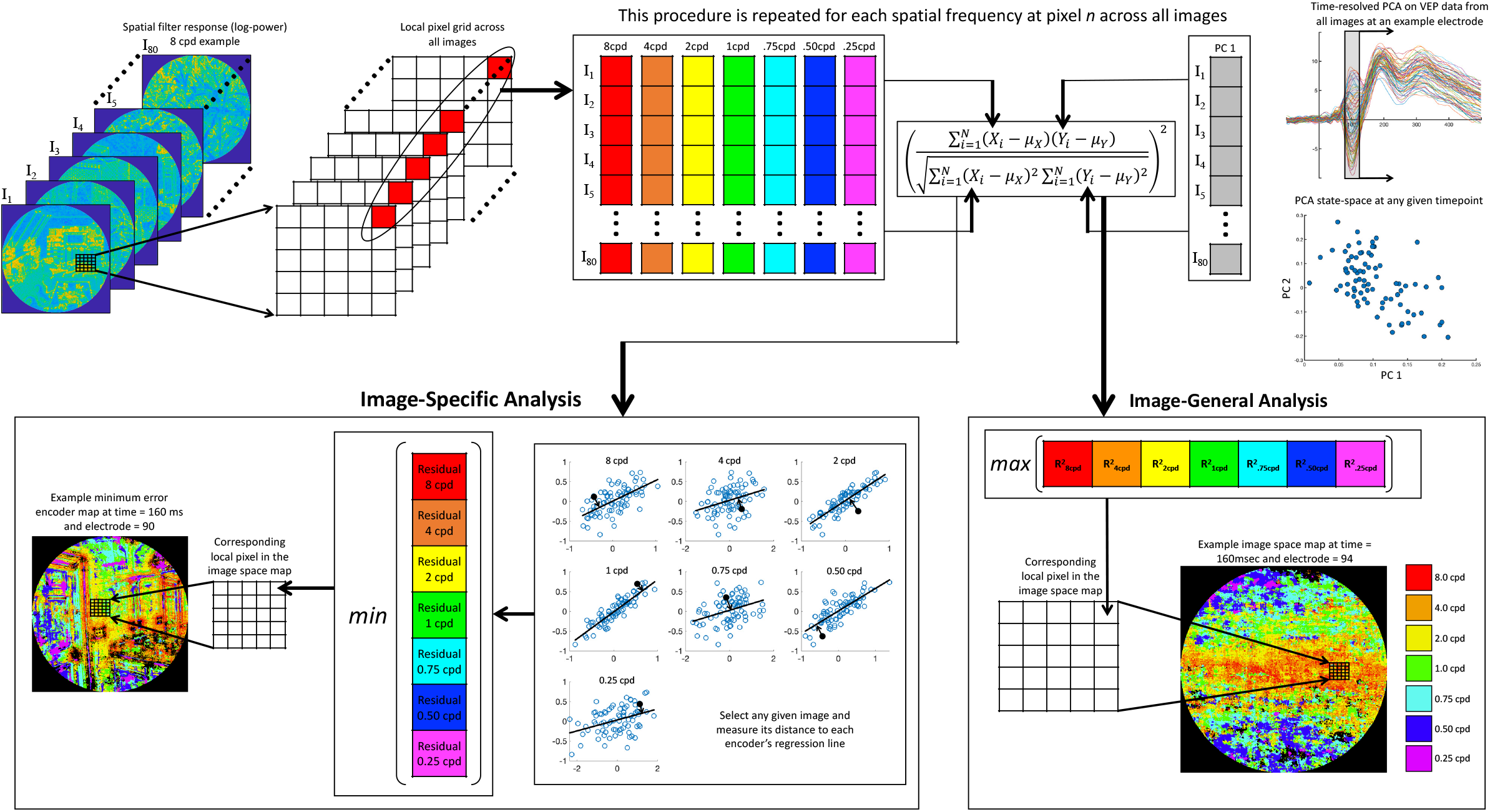
Schematic illustration of the DETI mapping procedure. All stimuli were represented in a log-Gabor filter power response space defined by seven peak SFs at each pixel coordinate across all stimuli (8 cpd encoder response examples in the upper left). For any given time point and electrode, and at any given pixel coordinate, we sampled the filter responses across all images for a given encoder and assembled them into an 80 × 1 array (this was repeated for each encoder). Then, for any given time point and electrode, each encoder’s 80 × 1 filter response array was regressed against the first PC taken from VEP data evoked by all images at the corresponding time point. Two types of maps were then produced: image-general maps (bottom right) and image-specific maps (bottom left). The image-general tagged each pixel coordinate with the SF of the encoder that had the highest R^2^. Similarly, the image-specific analysis was designed to find the encoder that could best predict the VEP variability across all images captured by the first PC for any given stimulus. However, instead of using the best fit across all images, this analysis finds the encoder regression fit that had the shortest distance (i.e., smallest residual) to a given image and tagged the corresponding pixel location with that encoder’s SF.

### Time-resolved dimension reduction of VEP data

We recorded VEP data from human participants (n = 24) while they viewed 80 scene images sampled from a variety of environments. We focused our analyses on the posterior scalp electrodes (54 in total; see **Supplementary Figure 1**) because VEPs recorded at those sites carry retinotopically selective spatial frequency (SF) information [36]. To reduce the high-dimensionality of each participant’s VEP data (54 electrodes X 500 time points), we applied time-resolved PCA at each electrode across all scene evoked VEPs and time points in steps of 5 ms, centered within a 41 ms temporal window (±20 ms from a given time point). Across all participants and time points, the first two PCs were found to explain 93.2% (median) of the VEP variance, with PC 1 accounting for 76.8% of the variance. The first PC’s eigenvector for each electrode and time point was therefore used to define a uni-dimensional ‘space’ that could be mapped to each scene location within a representational space defined by our log-Gabor encoder model.

### Log-Gabor Encoder Model

We used log-Gabor filter response power as a model of the response variability of differently tuned neurons at each location in our stimuli [37] (see the Materials & Methods section for more detail). Briefly, the model consisted of seven filters, each tuned to a different peak spatial frequency (SF) (0.25, 0.50, 0.75, 1, 2, 4, or 8 cycles per degree; cpd) and all orientations (i.e., a log ‘doughnut’ filter in the Fourier domain) – refer to **Figure 3** for examples. The code to build the encoders and to generate the encoder space can be downloaded here [<u>**LINK**</u>]. While the focus of this study was on SF, we nevertheless built a set of filter encoders that spanned all SFs centered on eight different orientations (0 – 157.5° in steps of 22.5°) and report some of those mapping results as Supplementary Material.

### Linking VEP Data to Encoder Space and DETI Mapping Procedures

The linking operation begins by mapping each pixel coordinate across a set of scenes to each encoder’s peak SF. Thus, each pixel coordinate has seven dimensions, each defined by filter response magnitudes across all images in our stimulus set. Each filter’s response at a given pixel coordinate across all images is then assembled into an array (illustrated in **Figure 3**). Next, each filter’s pixel array is regressed against the first PC’s eigenvector from the VEP data at each 5 ms time step and electrode. This procedure creates seven 2D encoder maps for each electrode and time point, with each cell in a given map containing an R^2^ value (see **Figure 4A** for example encoder R^2^ maps).

**Figure 4.**
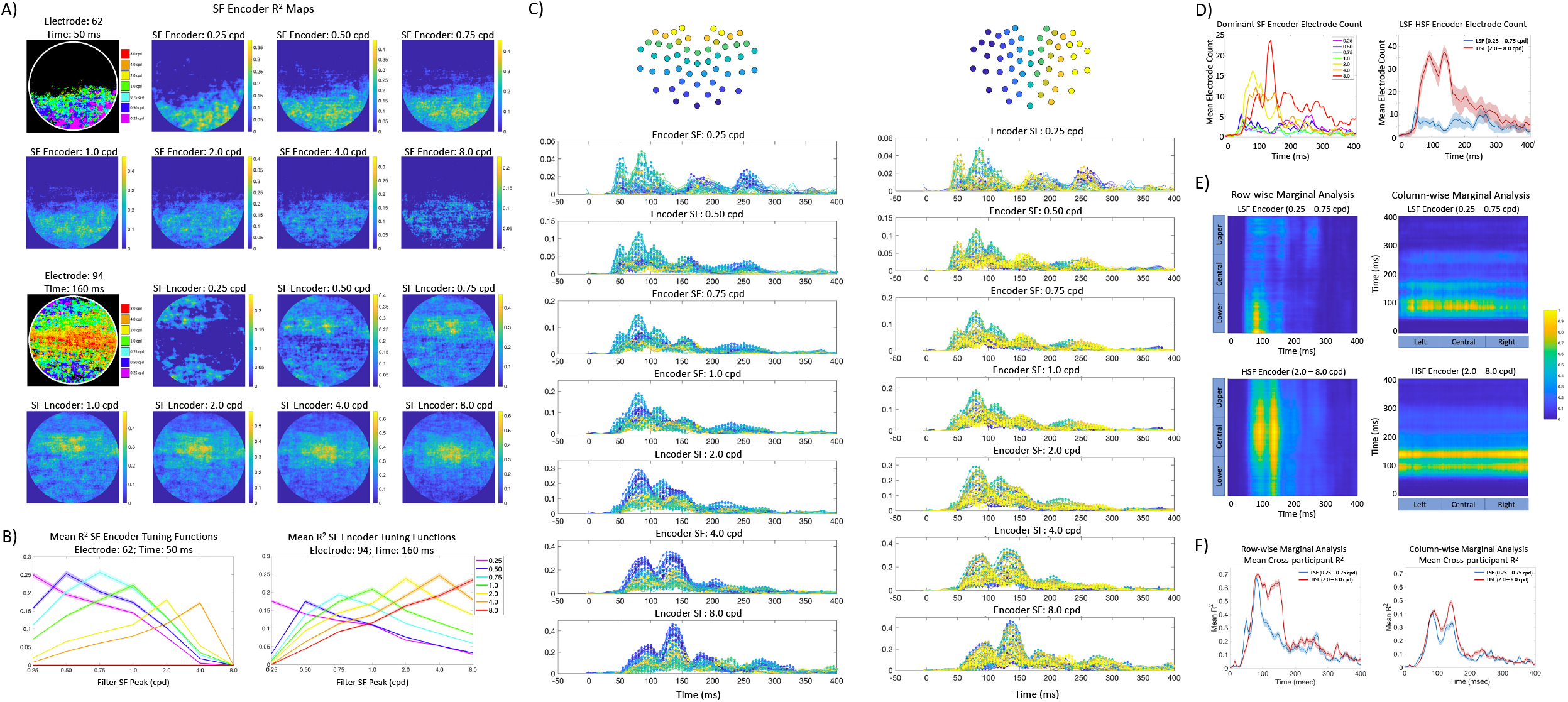
Image-general DETI mapping procedure results. **A)** Example R^2^ maps from two different electrodes and time points. The DETI maps for each example are shown in the upper left of each set of R^2^ maps. Each R^2^ map shows significant R^2^s at each pixel location in image space. The colorbar for each map shows R^2^. **B)** Example encoder R^2^ tuning functions for the two DETI maps shown in (A), averaged over all instances of each encoder’s tag in the DETI maps (y-axis is averaged R^2^, x-axis is encoder peak SF). The shaded region of each trace shows the 95% confidence interval over all instances of pixels for each encoder. **C)** Encoder probability over time, averaged across participants. The y-axes show participant-averaged encoder probability (note that the axes are different across SF peak), with time (ms) on the x-axes. Each trace is from a specific electrode. Probabilities that were above the lower bound of 95% confidence intervals across participants are indicated with a marker point. The electrode traces are color-coded topographically in two ways (illustrated at the top of each set of plots). The left-hand plots are coded from ventral-posterior to dorsal-posterior portions of the scalp, with the right-hand side coded from left to right across the scalp. **D)** Participant averaged electrode counts based on encoder dominated DETI maps. The left-hand plot shows the number of electrodes that were dominated (largest sum) by each of the seven encoders. The right-hand side is a summary of that plot and shows the electrode counts summed over LSFs (0.25 - 0.75 cpd) and HSFs (2.0 - 8.0 cpd). The shaded area shows the 95% confidence intervals over participants. **E)** Marginal analyses of the 2D DETI map histograms for the lower (LSF) and higher (HSF) encoder peak SF. The row-wise marginal analysis consists of an average across the R^2^ maps from left to right for each time point. The columns of that plot were first normalized over time and then normalized again within each column to emphasize encoder fit magnitude over time and space. The column-wise marginal analysis was carried out the same way, but from top to bottom of the R^2^ maps (the normalization therefore took place row-wise). For a dynamic view of each marginal analysis, follow this link [<u>**LINK**</u>]. **F)** Averaged cross-participant R^2^s for each column in the row-wise marginal analysis (left) and each row of the column-wise marginal analysis (right). The shaded region shows the 95% confidence interval across participants.

At this point, the analysis splits to form two types of mapping procedures: 1) an image-general spatiotemporal view of scene SF coding over the entire stimulus set, and 2) a scene-specific spatiotemporal view of the SF code for each scene in our stimulus set. Specifically, the first approach builds maps based on local (i.e., pixel level) encoder variability across all stimuli at each time point (**Figure 3** lower right). That is, each pixel is ‘tagged’ with the encoder’s peak SF that best predicted the first PC’s eigenvector. The second approach finds the encoder that best predicts the first PC at each pixel coordinate within each stimulus, thereby providing scene specific maps over time (**Figure 3** lower left). Specifically, each pixel coordinate for a given scene is tagged with an encoder’s peak SF based on the shortest distance (smallest residual) between that scene and any one of the encoder regression lines across all images.

### DETI Image-General Mapping Analysis & Results

Example DETI maps from participant averaged data are shown in **Figure 1**. Please view the accompanying movie for the complete depiction of how different DETI maps evolve over time [<u>**LINK**</u>]. Prior to building the image-general DETI maps, all encoder R^2^ maps (**Figure 4A**) were corrected for multiple comparisons using the Benjamini-Hochberg procedure with a false discovery rate of 5%. We verified the validity of the DETI mapping procedure by running it on participant averaged trial shuffled data. The resulting Benjamini-Hochberg corrected shuffled data maps over all electrodes at each time point were either completely empty (~94.6% over all electrodes and time) or contained ~0.93% (median) pixels with encoder tags, demonstrating that this mapping procedure is largely resistant to noise. The Benjamini-Hochberg correction remained robust to false positives out to a false discovery rate of 10% (~3.32% pixels with encoder tags).

**Figure 4A** shows example DETI maps for two electrodes at two different time points, alongside the encoder R^2^ maps that were used to tag the image-general maps based on the encoder’s peak SF that yielded the largest significant R^2^. Selecting the maximum R^2^ assumes that there is an underlying tuning function that drives the R^2^s for each encoder. **Figure 4B** verifies this assumption by showing each encoder’s R^2^ averaged across all instances of its tag in the corresponding DETI maps and shows that each encoder exhibits an R^2^ tuning function measured over the seven peak SFs of our encoder model (see **Supplementary Figure 2** for a comprehensive analysis).

The DETI procedure enabled us to estimate the general coding principles of scenes over time. The time-resolved maps shown in **Figure 1** (and the corresponding movie) reveal differences with respect to 1) scalp topography, 2) the relative contribution of SF encoders to each DETI map, and 3) how the encoders map to general image regions over time. To quantify those observations, we reduced the dimensionality of the DETI maps in several ways. First, we calculated the probability of observing pixels tagged with any given encoder’s peak SF by summing the number of pixels tagged by each encoder for each electrode at each time point and then divided each sum by the total number of visible pixels in the stimuli. The resulting encoder probability-by-time calculations for each electrode were then averaged across participants and shown in **Figure 4C** (see **Supplementary Figure 3** for the results from the orientation analysis). To investigate the extent to which any given encoder dominated the DETI maps across the scalp over time, we tagged each electrode with the encoder SF that was most prevalent in its map and summed the number of electrodes dominated by each encoder at each time point (**Figure 4D**).

To assess how the encoders map to general image regions over time, we analyzed the 2D map variation over all electrodes at each time point. Because cortical folding varies from person-to-person (which influences how VEP signals register on the scalp), we could not simply collapse across participants at corresponding electrodes. Instead, we converted each electrode’s fully tagged DETI map to a set of seven binary maps, one for each encoder, and then summed those maps (pixel-by-pixel) across all electrodes at each time point. The resulting 2D histograms for each encoder were then summed across participants. To provide a comprehensive view of the spatial biases in encoder maps over time, we first grouped the lower SF encoders (LSF; 0.25 - 0.75 cpd) and higher SF encoders (HSF; 2.0 - 8.0 cpd) and conducted an upper-to-lower and left-to-right marginal analysis on the LSF and HSF maps at each time point (**Figure 4E**). Finally, to assess participant agreement between the different marginal analyses, we regressed the upper/lower and left/right biases at each time point across participants (**Figure 4F**).

### DETI Image-Specific Mapping Analysis & Results

Example image-specific DETI maps from participant averaged data are shown in **Figure 2**. Please view the accompanying movie for the complete depiction of how different DETI maps evolve over time [<u>**LINK**</u>]. Additional image-specific map examples are shown in **Figure 5**. All image-specific mapping was based on the regression fitting procedure that was carried out for the image-general analysis, but here was focused on mapping based on minimal residual error from each encoder’s regression line. The trial shuffled analysis reported above suggests that the Benjamini-Hochberg correction procedure tended to be overly conservative, so we adjusted the false discovery rate to 10% for the analyses reported here (the results from analyses with a 5% false discovery rate were largely consistent with those reported here and are shown in **Supplementary Figure 4**).

**Figure 5.**
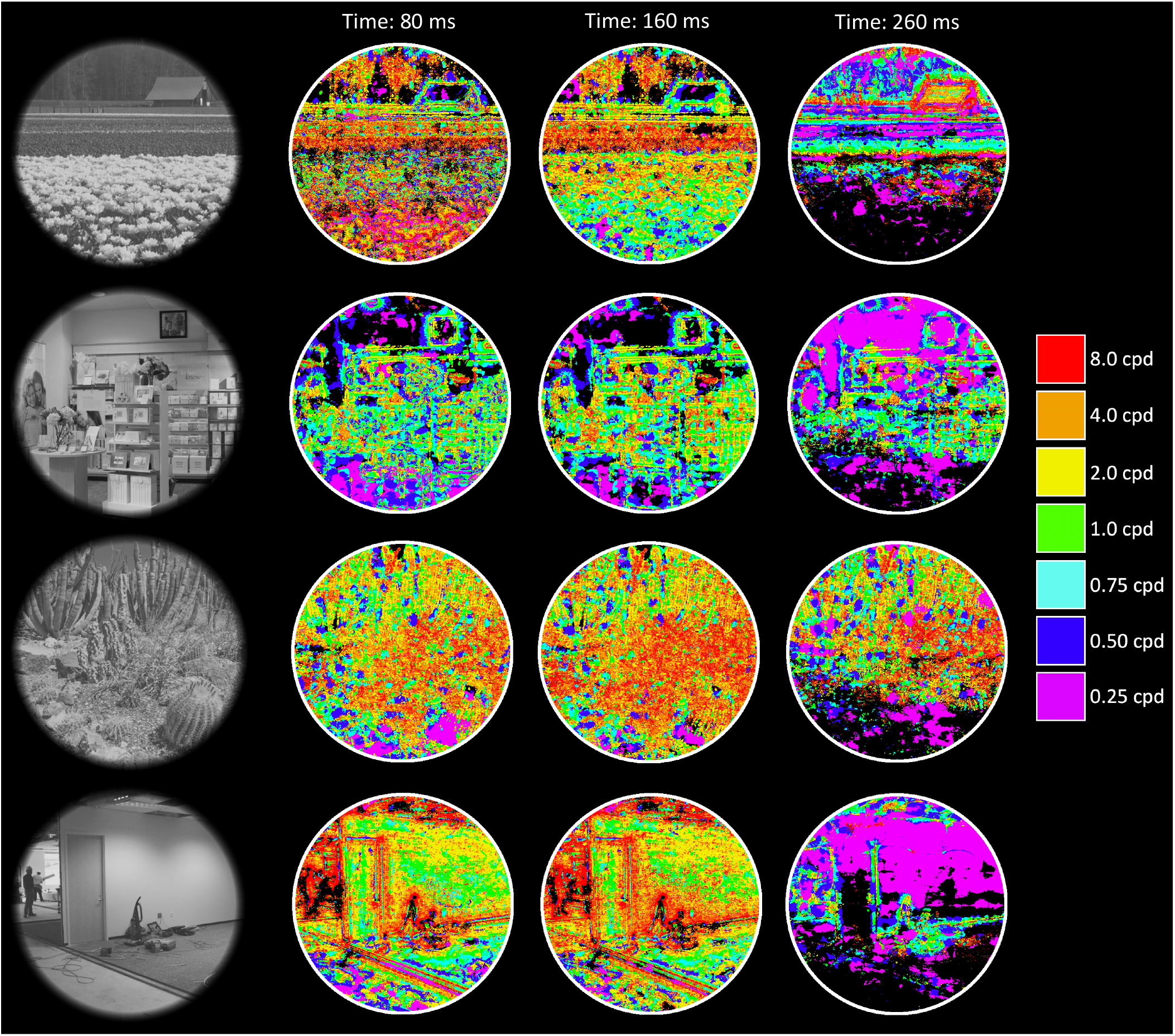
Additional stimulus examples and their DETI maps at different time points from electrode 90.

Together, **Figures 2** and **5** (and the corresponding movie) illustrate some of the diversity of image transformations over time. While the image-general analysis revealed the general coding principles across image space for our set of stimuli, this particular mapping procedure allowed us to characterize each image and image location according to the relative contributions of different encoders over time. As in the image-general analysis, we first calculated the probability of observing pixels tagged with any given encoder’s peak SF for each image and participant, which was then averaged across all participants. Example probabilities for each encoder over time for two example images are shown in **Figure 6A**. With the encoder probability data in hand, we were able to project all images into a state-space representation based on all electrodes at each time point for each participant. That particular analysis is illustrated in **Figure 7**. Briefly, for any given time point, each encoder’s probability for each electrode was stored in an array on an image-by-image basis, resulting in a 378 × 80 matrix for each time point (i.e., 54 electrodes * 7 encoders X 80 scenes). Thus, each image is represented as patterns of SF probability across encoders and electrodes. Including all 54 electrodes allowed representations based on visual signals from different regions of the visual field [36,38]. The resulting matrix was then submitted to PCA. The first two PCs accounted for 87.4% (median) of the variance across time, images, and participants, with the first PC accounting for 80.7% (median). All images could then be represented in an SF probability state-space defined by the first two PCs at each time point and participant (examples of that space for a given participant at three different time points are shown in **Figure 6B**).

**Figure 6.**
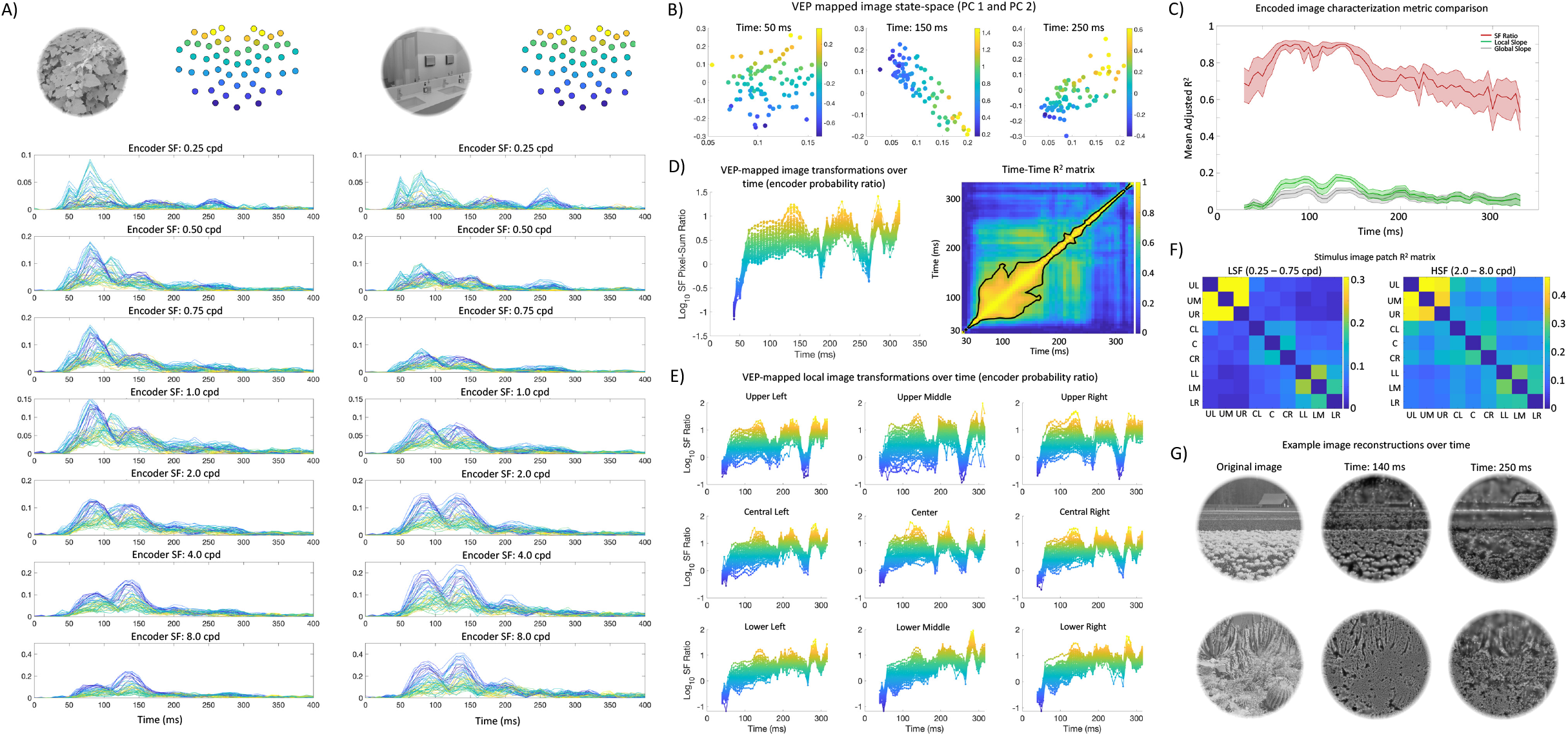
Image-specific DETI mapping procedure results. **A)** Encoder probability over time for two stimulus examples (upper left of each set of plots), averaged across participants. The y-axes show participant-averaged encoder probability, with time (ms) on the x-axes. Each trace is from a specific electrode and color-coded from ventral-posterior to dorsal posterior electrode (illustrated at the upper right of each set of plots). **B)** Principal component (PC) defined VEP-mapped image state-space from an example participant at three different time points (refer to text and **Figure 7** for further details). Data points are images, and the colorbar corresponds to the log10 HSF-to-LSF probability ratio for each image in that space. The x- and y-axes are PC 1 and 2 respectively. **C)** Characterization metric comparison for explaining the relative location of each image in PC-defined state-space over time. The y-axis shows the average adjusted R^2^ across participants. The shaded region shows the 95% confidence interval of the fits across participants. **D)** The left-hand plot shows participant averaged log10 HSF-to-LSF probability ratio for each encoder tagged image over time across the entire stimulus space (i.e., all pixels). On the right-hand side is the participant averaged time-time R^2^ matrix assembled by regressing the log10 HSF-to-LSF probability ratios across images for each time point against every other time point (colorbar is R^2^). The contour line encapsulates the top 12% of the R^2^s. **E)** Participant averaged log10 HSF-to-LSF probability ratio for each encoder tagged image over time, measured at nine different locations within stimulus space (see text for more details). The format of the plots is the same as in (D, left). **F)** R^2^ matrices showing the relationship between the encoder responses at each pixel coordinate within one of the nine image regions and the corresponding pixel coordinate in every other patch region. R^2^s are averaged across all pixel coordinates within each patch, and then averaged across the lower SFs (LSF; left hand matrix) and higher SFs (HSF; right hand matrix). The colorbar shows R^2^. **G)** Example stimuli along with their SF filter reconstructions based on the 2D encoder histograms at two time points (see Materials & Methods for further details).

**Figure 7.**
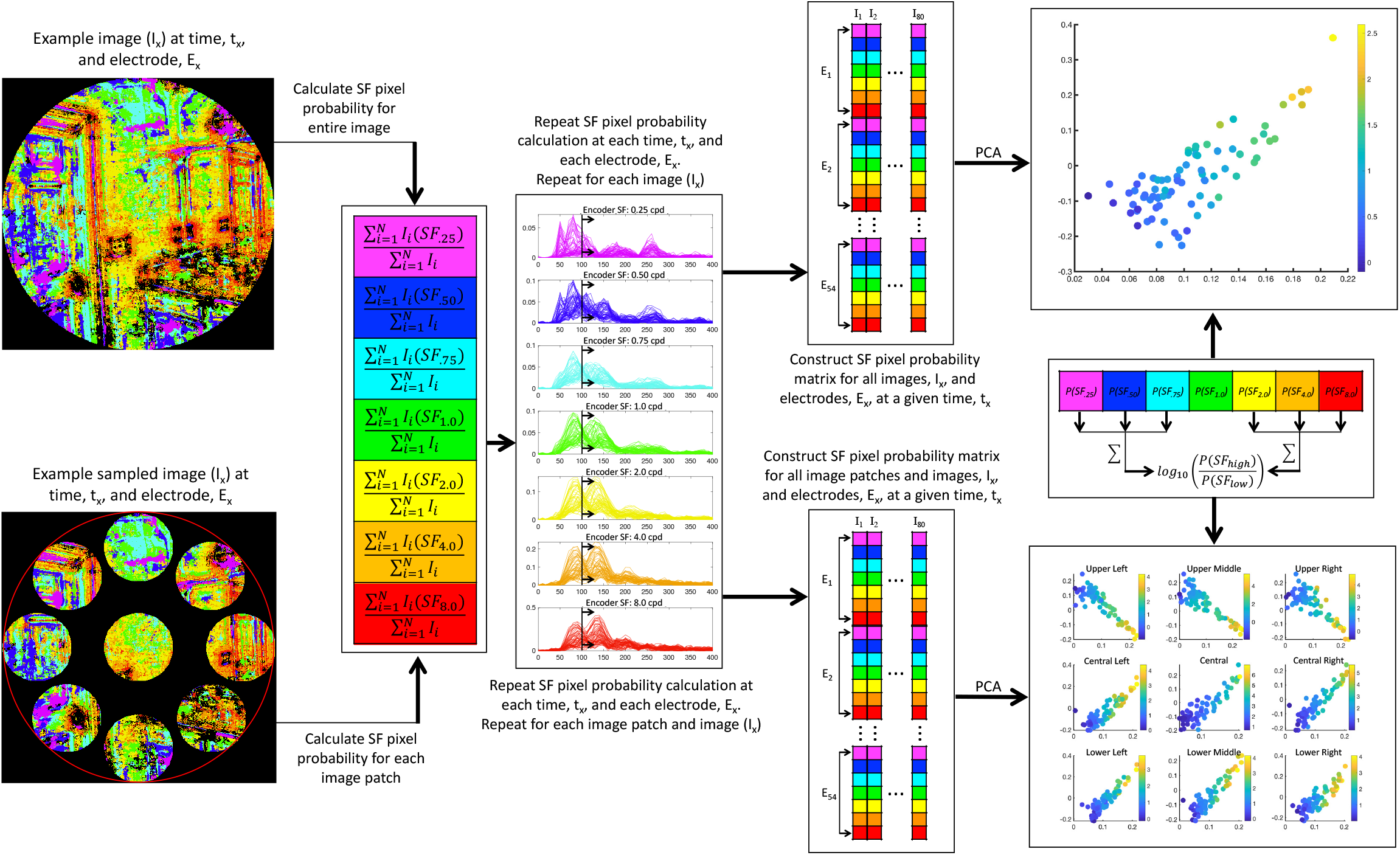
Illustration of the procedure for projecting all encoder-tagged images into a state-space representation at each time point for each participant using all electrodes. This procedure was either carried out for entire images (top left) or within patches located at nine different regions of the images (bottom left). Encoder probabilities were calculated as described in the text (and **Figure 4**) for whole images or image patches. At each time point and across all electrodes, each encoder’s probability was stored in a matrix as an array on an image-by-image basis (e.g., a 378 × 80 matrix for each time point). Each matrix was then submitted to PCA, with the first two PCs defining the encoded image’s state-space. Each image was then characterized by taking the log10 HSF-to-LSF probability ratio (see text for further detail).

Next, we sought to characterize the relative positioning of the images in that space according to encoder probability. Specifically, we characterized each image by the ratio of the summed HSF probabilities (2.0 – 8.0 cpd) to the summed LSF (0.25 – 0.75 cpd) probabilities at each time point as depicted in **Figure 7**. To assess the ability of this characterization to explain the first two PCs, we submitted the SF ratio for each image (and at each time point) to multiple regression and show the adjusted R^2^ over time in **Figure 6C**. We also compared this characterization to that which could be explained by simple Fourier image statistics based on: 1) the amplitude spectrum slope of each image, and 2) the ratio of local filter response slopes (shallow slopes to steep slopes) at each pixel (also shown in **Figure 6C**). The success of the encoder SF ratio characterization of images in SF probability PC space enabled us to visualize each image’s transformation over time via the ratio characterization (participant averaged ratios: **Figure 6D** left). As a final analysis, we ran a time-time regression analysis on the participant averaged SF probability ratios to observe the temporal generalization of the image transformations over time (**Figure 6D** right).

In order to better understand the unique transformational states observed across images (as defined by the SF ratio characterization), we assessed the transformations within localized image windows. Out of practicality, we ran this analysis within nine different spatial windows using a polar coordinate system to define the position of each spatial window (refer to **Figure 7** for an example). Each spatial window had a diameter of 128 pixels (4.7° of visual angle). The center-to-center distance between the center window and the outer windows was 190 pixels (7.1° of visual angle), with the center-to-center distance between each outer window being 160 pixels (6° of visual angle). The analysis described above to produce **Figure 6D** (left) was carried out on each window location across all images on a participant-by-participant basis (**Figure 6E**). To verify that the local differences in transformational states are a reflection of unique encoder information contained within those regions, we regressed the encoder responses at corresponding pixel locations between the spatial windows for each image, and then averaged the pixel R^2^s within each window, and then across images. The resulting window-wise R^2^ matrices (averaged over 0.25 – 0.75 cpd for LSF, and 2.0 – 8.0 cpd for HSF) are shown in **Figure 6F** (the same analysis was applied to all images in our database and yielded virtually identical results). Finally, to provide a rough estimate of how the images may appear in different transformational states, we ‘reconstructed’ example images using an approach that is similar to the SF bubbles technique in the spatial domain [39] (see Materials & Methods for details). The examples shown in **Figure 6G** show two time points, with an earlier time point (140 ms) showing the reasonably constant SF mapping across image region, with a later (250 ms) inconsistent representation across image region where the upper portion of the images are in an LSF transformational state and lower portion of the images are in an HSF transformational state.

## Discussion

### Image-general DETI Mapping

The encoder probability-by-time analysis (**Figure 4C**) shows that the posterior electrodes vary with respect to where on the scalp that the dominant encoder SF is expressed, likely reflecting their underlying calcarine sources in accordance with the cruciform model [38]. Specifically, HSFs tend to dominate the DETI maps along the ventral-posterior electrodes, with LSFs dominating the DETI maps along the dorsal-posterior electrodes – both observations are consistent with previous literature [40]. This consistency shows that the DETI procedure can map the SF scalp topography without resorting to multiple experiments that present stimuli at different locations in the visual field to avoid dipole cancellation. No clear lateralization was observed (**Figure 4C**, right hand side). Further, the analysis focused on SF encoder bias over electrodes at each time point (**Figure 4D**) shows that LSFs dominate early (~50 ms), followed by the HSFs, with the 2 cpd and 4 cpd encoders preceding the 8 cpd encoder. Interestingly, every time point yields electrodes that are dominated by different encoder SFs, revealing a multiscale representation of scenes over time. The results from the orientation-based encoders do not show a clear topographic organization, but instead show that orientations at and near horizontal have the lowest probability in the maps over time (**Supplementary Figure 3**). Such a result is consistent with normalization operations in visual cortex and scene selective cortices that operate to reduce the magnitude of horizontal information [41–42].

Regarding the image-general regional biases that the DETI mapping procedure uncovered, the 2D histogram analysis (**Figure 4E**) shows that the upper and lower portions of images are coded by LSFs, with the HSFs coding the central portion of images. These biases were largely consistent across participants, especially during the earlier time window (e.g., 50 – 180 ms) (**Figure 4F**). Interestingly, those spatial biases were not static, but varied in a nonuniform manner over time. Specifically, the lower portion of image space is dominantly coded early (~50 ms) by LSFs, with the upper portion of image space dominating later (~250 ms), again with LSFs. The central portion of image space shows two waves of HSFs, with the first wave more centralized than the second. When compared to **Figure 4D**, the first wave corresponds to 2 – 4 cpd, with the spatially broader wave corresponding to 8 cpd. Additional asymmetries were observed laterally across images with LSFs showing an early (~70 ms) central-to-left hand bias, with HSFs showing a slightly later (~80 ms) bias toward the far left and right portions of images. Interestingly, such nonuniformities over time would not be expected from a simple linear model based on retinotopic mapping of SF preference and suggest a differential emphasis on image regions over time, thereby providing insight into a possible prioritization of different image regions as time advances. For instance, an early prioritization of the ground plane may support rapid judgements regarding scene navigation [43–44], with a later upper image region analyses focused on landmark organization [45].

### Image-specific DETI Mapping

Consistent with the image-general mapping, the encoder probability-by-time analysis (**Figure 6A**) shows that individual images exhibit a tendency for HSFs to dominate the electrode maps along the ventral-posterior electrodes, with LSFs dominating the electrode maps along the dorsal-posterior electrodes. Interestingly, each image showed a relatively unique encoder-tag probability over time and electrode. The unique tag probability over time enabled us to characterize each image’s transformational state at any given time point with the SF probability ratio. The regression analysis reported in **Figure 6C** shows that the SF probability ratio characterization provides an excellent account of the relative positioning of the stimuli in PC-defined SF probability state-space (i.e., **Figure 6B** top) for all participants. Further, that characterization is far superior to simple image statistic characterizations, meaning that the different transformational states that the images undergo are unlikely to be explained by simple Fourier image statistics.

Of particular interest was the analysis of different image transformational states over time (**Figure 6D** left). Here, the DETI mapping revealed an interesting, possibly two-stage [46], pattern of representations in that the images first show an initial LSF based code (~50 ms), followed by a relative HSF code (~70 ms to ~140 ms) where the encoder SF ratio variance is ~3 times larger than any other time window. Around 150 ms, the image transformations undergo what appears to be intermittent LSF transformations at ~180 ms and ~260 ms, possibly indicative of recurrent processes [47–48]. The temporal comparison of the transformation states of the images (**Figure 6D** right) shows an increase of temporal variability (which can be interpreted as a reduction in temporal similarity) between representational states starting at ~240 ms, suggesting that each transformation becomes more unique as time advances.

Interestingly, the results from the local window analysis (**Figure 6E**) reveal that different locations within image space undergo a relatively unique transformational process over time, with the largest difference being between the upper and lower portions of image space, with the central portion of image space being somewhat intermediate. Specifically, the upper and central portions of different images show similar transformational states over time compared to the entire image (**Figure 6D** left), with the lower portion of an image showing transformational states that advance from LSF based representation to HSF based representations. A time-time R^2^ analysis on each location revealed results that were somewhat similar to the full image analysis, with the upper and central portions of image space showing the gradual reduction of similarity between representational states over time. On the other hand, the lower portion shows overall less similarity between the representational states (see **Figure 8**). This suggests that the transformational states are more unique from time point to time point in the lower portion of image space compared to the upper portion. What’s interesting about this differential location coding is that the analysis on encoder similarity (**Figure 6F**) shows that the encoder information contained *within* each region (e.g., upper, central, and lower) is more similar than when compared to the information contained within the other two regions, further supporting differential neural coding operation at different locations within images over time. Importantly, those differential transformational states within different image regions suggest that the temporal coding of visual information is far more complex than a simple coarse-to-fine analysis and subsequent mapping to higher cortical representations [49–50].

**Figure 8.**
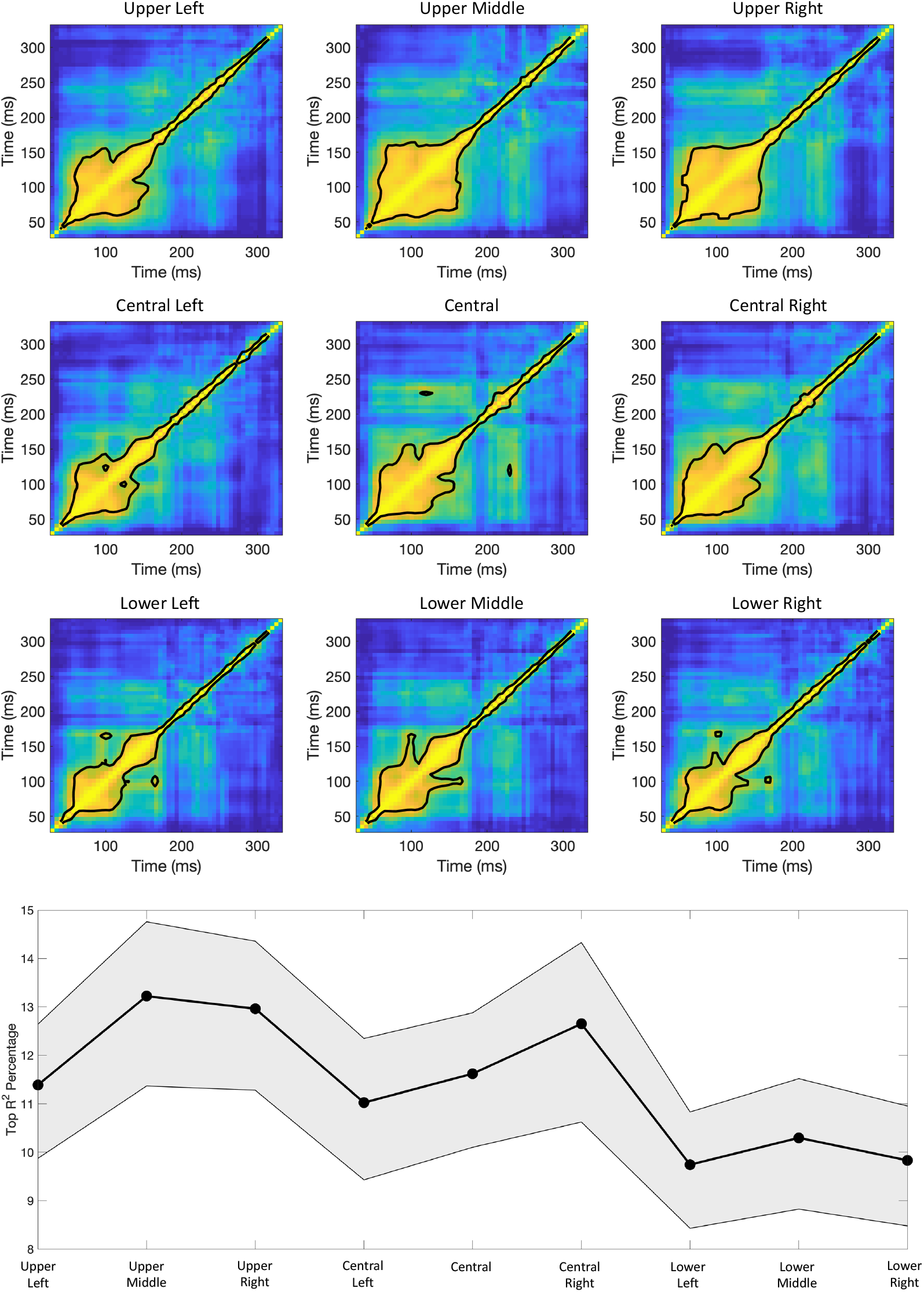
Top: Results from the time-time regression analysis on the participant averaged SF probability ratios within each of the nine local image windows. Bottom: The percentage of time-time R^2^s that fell within the top 12% contour area was calculated for each participant, and then averaged across participants (gray area shows the 95% confidence interval). That plot illustrates the tendency of the lower image windows to result in more temporal variation, thereby suggesting more unique temporal SF ratio characterizations on a time point by time point basis. This was first verified with a one-way ANOVA F(8,207) = 2.9, P = .005, with post t-tests showing significant differences between all lower windows and the upper middle, upper right, and central right windows (P’s < .02).

### Concluding Remarks

The DETI mapping procedure offers many advantages over traditional EEG component measures by providing a framework to assess how each image region contributes to the underlying VEP signals. Specifically, electrodes over the posterior scalp tend to contain signals that carry information in the upper or lower peripheral visual fields, while electrodes over the occipital pole tend to signal for information in the central visual field [40]. However, using raw VEPs to assess the entirety of the visual signal simultaneously for large-field scene images yields overlapping components that differ in polarity as a function of visual field location. What that means is that the complete spatial representation of the visual signal would be largely obscured by dipole cancellation [38]. The DETI mapping procedure circumvents this problem by relying on low dimensional signal *variance* to map VEPs to scene images in encoder space on a pixel-by-pixel basis, thereby enabling a relatively clear visualization of the transformational states that images undergo. Further, DETI mapping provides a rich dataset (54 electrodes X 206,643 pixels X 7 encoders = 78,111,054-dimensional space per time point) for a variety of characterizations, thereby enabling multiple levels of downstream mapping of that code. For example, the relative transformational states between scenes and scene regions can be used to project DETI maps into a high-dimensional state-space for representational similarity analysis (RSA) at each time point. Such an analysis would allow insight into how different transformational states of scene regions map onto the knowledge structures that drive intelligent behavior [11–15]. Further, DETI mapping is an encoding based approach, meaning that it can be easily outfitted with encoding models from other sensory modalities or multiple non-linear encoders within a single modality [26]. The DETI procedure can also accommodate ‘higher level’ encoder models derived from human labeling of scenes. Such an approach would enable the mapping of neural activity to image regions linked to task relevant information, thereby providing an opportunity to understand how early transformational states map onto later categorical representations [21] – a possibility that we are currently exploring. As powerful as the DETI mapping procedure is, it is not without its limitations. Because the procedure maps VEP variance to image variance in encoder space, any covariation between image regions will result in identical tagging, thus complicating any assessment seeking to disambiguate those regions (though sphering the encoder space may help in that regard [21]). Nevertheless, as demonstrated in the present study, DETI mapping holds much potential to advance our understanding of the spatiotemporal coding of visual information.

## Materials & Methods

### Apparatus

All stimuli were presented on a 23.6” VIEWPixx/EEG scanning LED-backlight LCD monitor with one ms black-to-white pixel response time. Maximum luminance output of the display was 100 cd/m^2^, with a frame rate of 120 Hz and resolution of 1920 × 1080 pixels. Single pixels subtended .0382° of visual angle as viewed from 35 cm. Head position was maintained with an Applied Science Laboratories (ASL) chin rest.

### Participants

A total of 35 participants were recruited for this experiment. Of those, 8 failed to complete both recording sessions and 3 were excluded for having fewer than 50% valid trials following artifact rejection. The age of the remaining 24 participants (13 female, 22 right-handed) ranged from 18-21 (median age = 18). All participants had normal (or corrected to normal) vision as determined by standard ETDRS acuity charts, gave Institutional Review Board-approved written informed consent before participating, and were compensated for their time.

### Stimuli

Stimuli were selected from a large database of real-world scenes consisting of 2500 photographs that varied in content from purely natural to purely carpentered (both indoor and outdoor), with various mixtures of natural/carpentered environments in between [37]. All images were 512 × 512 pixels and converted to grayscale using the standard weighted sum conversion in MatLab.

For the purposes of stimulus presentation and analysis, all images were calibrated according to the following procedures. First, each image was fit with a hard-edge circular window (with a diameter of 512 pixels) whereby all pixels that fell outside of the circular window were set to zero (i.e., we’re only interested in the 206,643 pixels that were presented to the participants as described later in this section). Next, each image was converted to an array, I(y), that included only the pixels that fell within the circular window and were made to possess the same root mean square (RMS) contrast and mean pixel luminance.

Root mean square contrast is defined as the standard deviation of all pixel luminance values divided by the mean of all pixel luminance values. Image arrays were set to have the same RMS contrast and zero mean using the following operations.

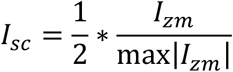

with I_zm_ defined as:

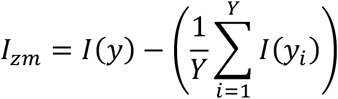

The pixels values of each image array are first normalized to fall between [−.5 .5] with zero mean as follows,

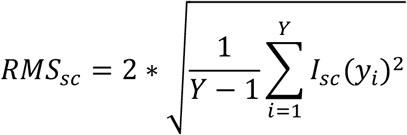

We then calculated an RMS scaling factor, S_rms_ = (2*RMS_t_)/RMS_sc_, with RMS_t_ set to a reasonable target RMS value. By reasonable, we mean a value that did not result in significant (> 5%) clipping of the resulting pixel values. That value was 0.20 for the images used in the current study. Finally, each image array was scaled to have an RMS equal to RMS_t_ and reassign to I(y) as follows: I(y) = 127*(I_sc_*S_rms_). Note that scaling by 127 puts the scaled pixel values of I(y) back in the original range of I_zm_.

Stimulus images were selected according to an image state-space sampling procedure as follows. All images in the database were left in vector form after RMS normalization (described above). When in array form, each cell constitutes a coordinate in a high-dimensional image state-space where each coordinate takes on a pixel luminance value ranging from [−127, 127]. An angular distance matrix was then constructed by calculating the angular distance (in degrees) between each image array and every other image array as follows:

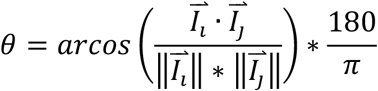

From there, the angular distance matrix was projected into a lower dimensional space (3D) via t-distributed stochastic neighbor embedding (t-SNE) [51]. Projecting the images into this space enables a general lower dimensional organization of images based on their structural attributes defined by pixel luminance.

Stimuli were selected by uniformly sampling 80 images from the lower dimensional t-SNE space in order to increase the probability of the different regions in our image state-space being represented in the stimulus set. All selected images maintained their RMS of .20 (defined above) but had their mean pixel luminance set to 127 and then fit with a circular linear edge-ramped window (512-pixel diameter, ramped to the mean pixel luminance) to obscure the square frame of the images. That step ensures that the contrast changes at the boundaries of the image were not biased to any particular orientation [52–53].

### Experimental Procedure

The experiment consisted of two recording sessions, each ranging between 50-55 min. Within each session, all 80 stimuli were presented 15 times, resulting in a total of 30 repetitions per image over both recording sessions (stimulus presentation order was randomized). Each trial began with a 500 ms fixation followed by a variable duration (500-750 ms) blank mean luminance screen to allow any fixation-driven activity to dissipate. The blank screen was immediately followed by the stimulus interval (500 ms) that was then followed by a variable 100-250 ms blank mean luminance screen, followed by a response screen. The response screen prompted the participant to rate the visual complexity of the stimulus image on a scale of 1-4 using a button box (response time was unlimited).

### EEG Recording and Processing

All Continuous EEGs were recorded in a Faraday chamber using Electrical Geodesics Incorporated’s (MagStim EGI) Geodesic EEG acquisition system (GES 400). All EEGs were obtained by means of Geodesic Hydrocel sensor nets consisting of a dense array of 128 channels (electrolytic sponges). The on-line reference was at the vertex (Cz), and the impedances were maintained below 50 kΩ (EGI amplifiers are high-impedance). All EEG signals were amplified and sampled at 1000 Hz. The digitized EEG waveforms were first highpass filtered at a 0.1 Hz cut-off frequency to remove the DC offset, and then lowpass filtered at a 45 Hz cutoff frequency to eliminate 60 Hz line noise.

Continuous EEGs were divided into 600 ms epochs (99 ms before stimulus onset and 500 ms of stimulus-driven response). Trials that contained eye movements or eye blinks during data epochs were excluded from analysis via magnitude thresholding followed by visual inspection. Additionally, all epochs were subjected to algorithmic artifact rejection whereby voltages exceeding +/− 100 μV or transients greater than +/− 100 μV were omitted from further analysis. These trial rejection routines resulted in a median of 9% (range 3% - 29%) of trials being rejected across participants. Each epoch was then re-referenced offline to the net average, and baseline-corrected to the last 99 ms of the blank interval that preceded the image interval. Finally, VEPs were constructed for each participant by averaging the processed epochs across trials for each image at each electrode, resulting in a 128 × 600 × 80 VEP data matrix for each participant. The full dataset (34Gb) can be downloaded here [<u>**LINK**</u>].

### Encoder Model Details

Visual evoked potentials constitute the sum of neural activity (post-synaptic potentials) at the circuit level. Given the retinotopic mapping of the visual cortices [34], the VEPs measured on the scalp likely stem from a summation of the underlying responses tuned to different image attributes at different image locations. If the majority of the summation arises from early visual cortical processes [1], then we can expect a good portion of the sum to be explained by contrast in different bands of spatial frequency and orientation. As a first approximation to model the relative response of differently tuned neurons at each location in our stimuli, we used a filter-power encoding model based on log-Gabor filters [37]. Specifically, the model consists of 7 filters, each tuned to a different peak spatial frequency (0.25, 0.50, 0.75, 1, 2, 4, 8 cpd) and all orientations (i.e., a log ‘doughnut’ filter in the Fourier domain). The spatial frequency bandwidths (full width at half height) of the filters scaled with peak spatial frequency such that they were broader at lower spatial frequencies and narrower at higher spatial frequencies: 2.3, 2.3, 2.0, 2.0, 1.75, 1.5, and 1.0 octaves respectively [2–3].

### Stimulus Representation in Encoder Space

All image filtering was conducted in the Fourier domain using the images in matrix form. To minimize edge effects in the Fourier domain due to the non-periodic nature of scene images, the images were symmetrized prior to taking the Fourier transform. Each symmetrized image was submitted to the 2D discrete fast Fourier transform to obtain H(u,v) as follows:

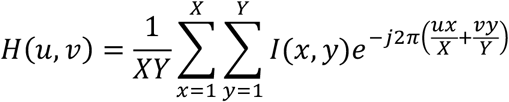

where I(x,y) represents a given image, with X and Y representing the dimensions of the symmetrized image. Next, the amplitude spectrum was calculated according to:

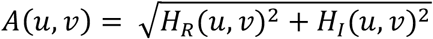

with H_R_(u,v) and H_I_(u,v) representing the real and imaginary parts of H(u,v), respectively. For filtering convenience, the amplitude spectrum, A(u,v) was shifted to polar coordinates and in this form will be denoted as A(*f*,θ), with *f* serving as the index along the radial (i.e., spatial frequency) dimension, and θ as the index along the theta (i.e., orientation) dimension.

Each image’s amplitude spectrum was then multiplied by a 2D log-Gabor filter. Log-Gabor filters in the Fourier domain consist of a log-Gaussian function along the *f* axis and a Gaussian function along the θ axis, which are then combined by multiplying a 2D log-Gaussian filter (i.e., a log ‘doughnut’ filter) with a 2D Gaussian ‘wedge’ filter. The construction of the 2D log-Gaussian filter, L_gaus_(*f*, θ), took place in the same polar coordinate frame as A(*f*,θ). Thus, for each θ axis, L_gaus_(*f*) was modulated as follows.

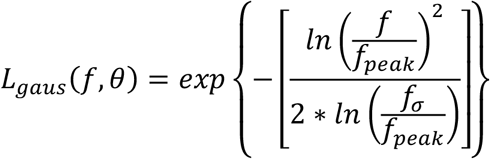

Where *f* increases with spatial frequency (radial distance), *f_peak_* represents the peak of the function, and *f_σ_* represents the SF bandwidth of the filter. Next, a 2D Gaussian function (modulated across θ in radians) about a central orientation was generated as follows.

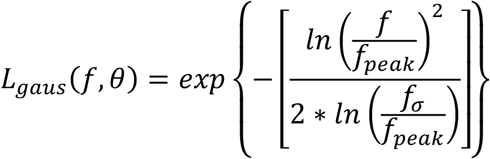

The log-Gabor filter, LG(*f*, θ), was then constructed by multiplying G_θ_(*f*, θ) by L_gaus_(*f*, θ).

The filtered amplitude spectra and corresponding phase spectra were then inverse Fourier transformed back into the spatial domain with the image in its original orientation cropped from the symmetrized version. We then took the natural log of the squared filter responses (i.e., each pixel location across all filters and images was expressed as log power). Representing the filtered images in the spatial domain allowed us access to the encoder responses at each pixel coordinate across all images. The code used to create the encoders and generate the encoder space can be downloaded here [<u>**LINK**</u>].

### Visualizing the transformational states of scenes over time

To provide a rough estimate of how the images may appear in different transformational states, we ‘reconstructed’ example images using an approach that is similar to the SF bubbles technique in the spatial domain [39]. Specifically, we created 2D encoder probability maps (summed over all electrodes) for a given image, then on a pixel-by-pixel basis, selected the encoder that had the highest electrode summed probability. We then sampled a window centered on the corresponding pixel of the image that had been filtered with the encoder at that SF. The diameter of that window scaled with encoder peak SF such that lower SFs had larger windows. Specifically, window diameter allowed for 1.5 periods of a given encoder’s SF. That sample was then weighted with a normalized Gaussian (normalized by area) and then summed with the corresponding pixel location in the reconstructed image template. This process was repeated for all pixels.

## Supplementary Material

**Supplementary Figure 1.**
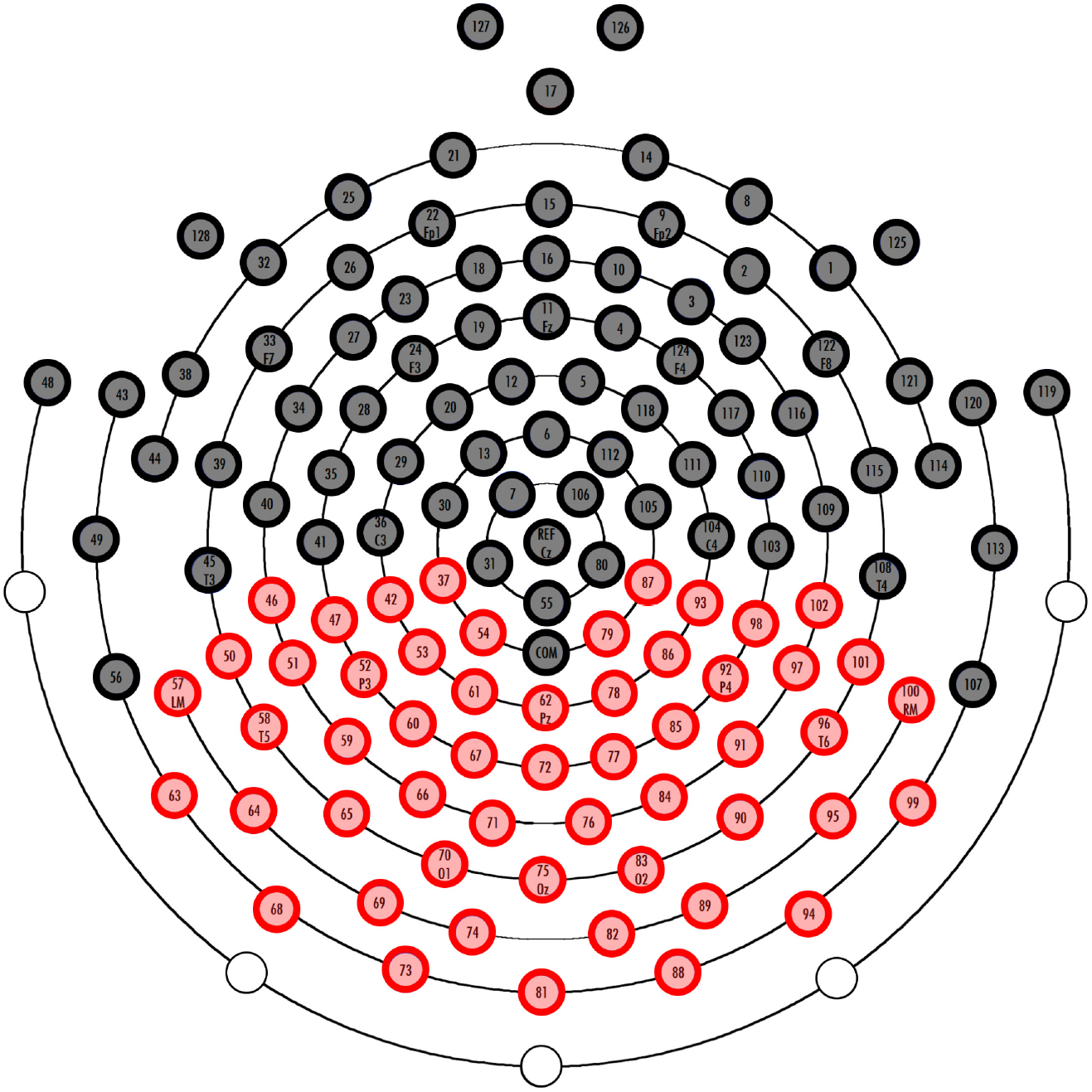
All EEG data were collected with Geodesic Hydrocel sensor nets consisting of a dense array of 128 channels. Above is the topographic representation of our sensor nets with the posterior electrodes that we included in our analysis highlighted in red. The posterior electrodes were chosen because VEPs recorded at those sites are known to carry retinotopically selective spatial frequency (SF) information.

**Supplementary Figure 2.**
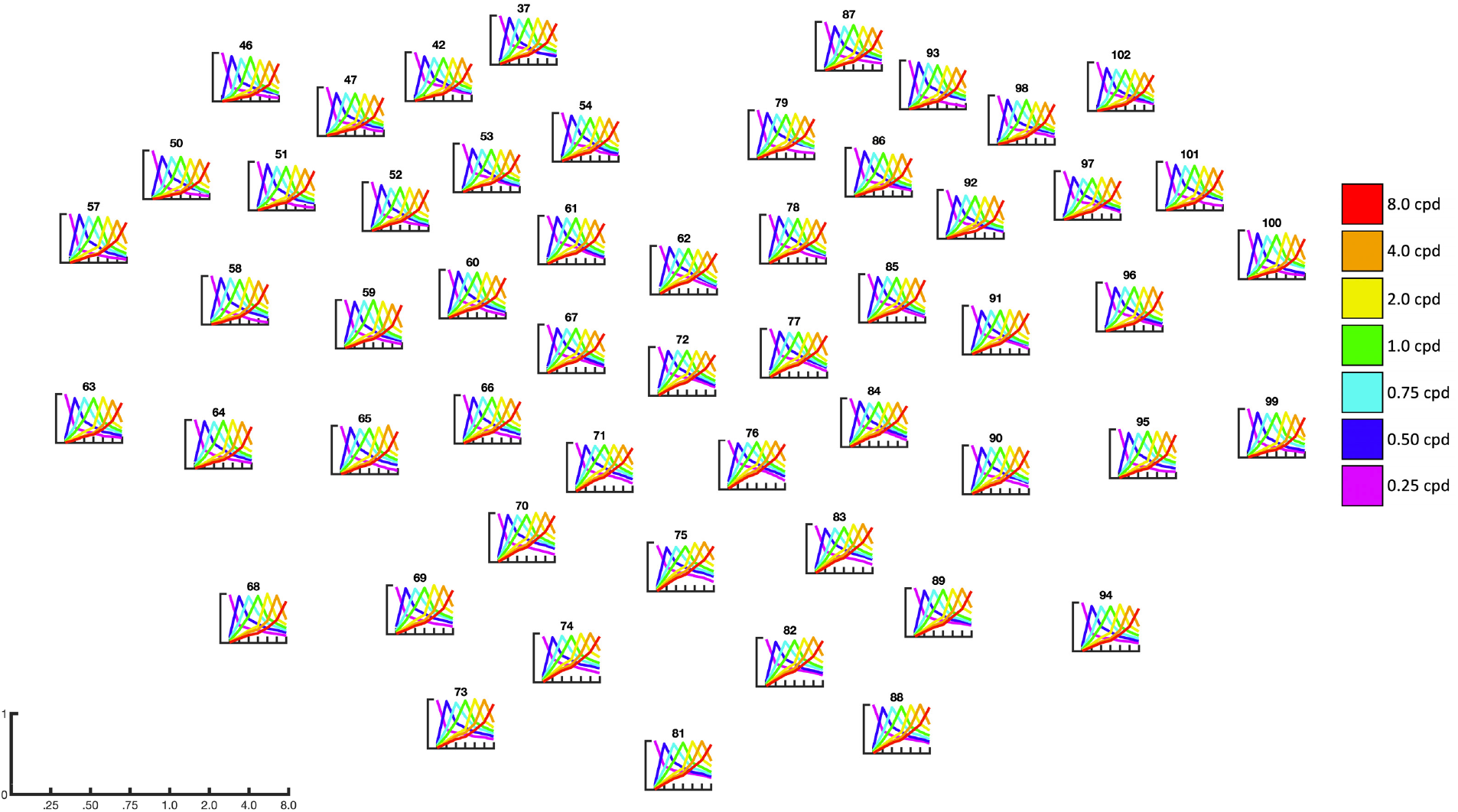
Posterior electrode plots showing each encoder’s R^2^ tuning function averaged over time. Specifically, we averaged across all instances of each encoder’s tag within each electrode’s DETI map at each time point for each participant, and then averaged across all time points and then across participants. All tuning functions have been normalized to the maximum peak within each plot (y-axis). The x-axis shows peak SF for each encoder (enlarged in the lower left corner). The results show largely similar tuning functions at each electrode, thereby justifying the use of selecting the largest R^2^ to tag each pixel in the DETI maps.

**Supplementary Figure 3.**
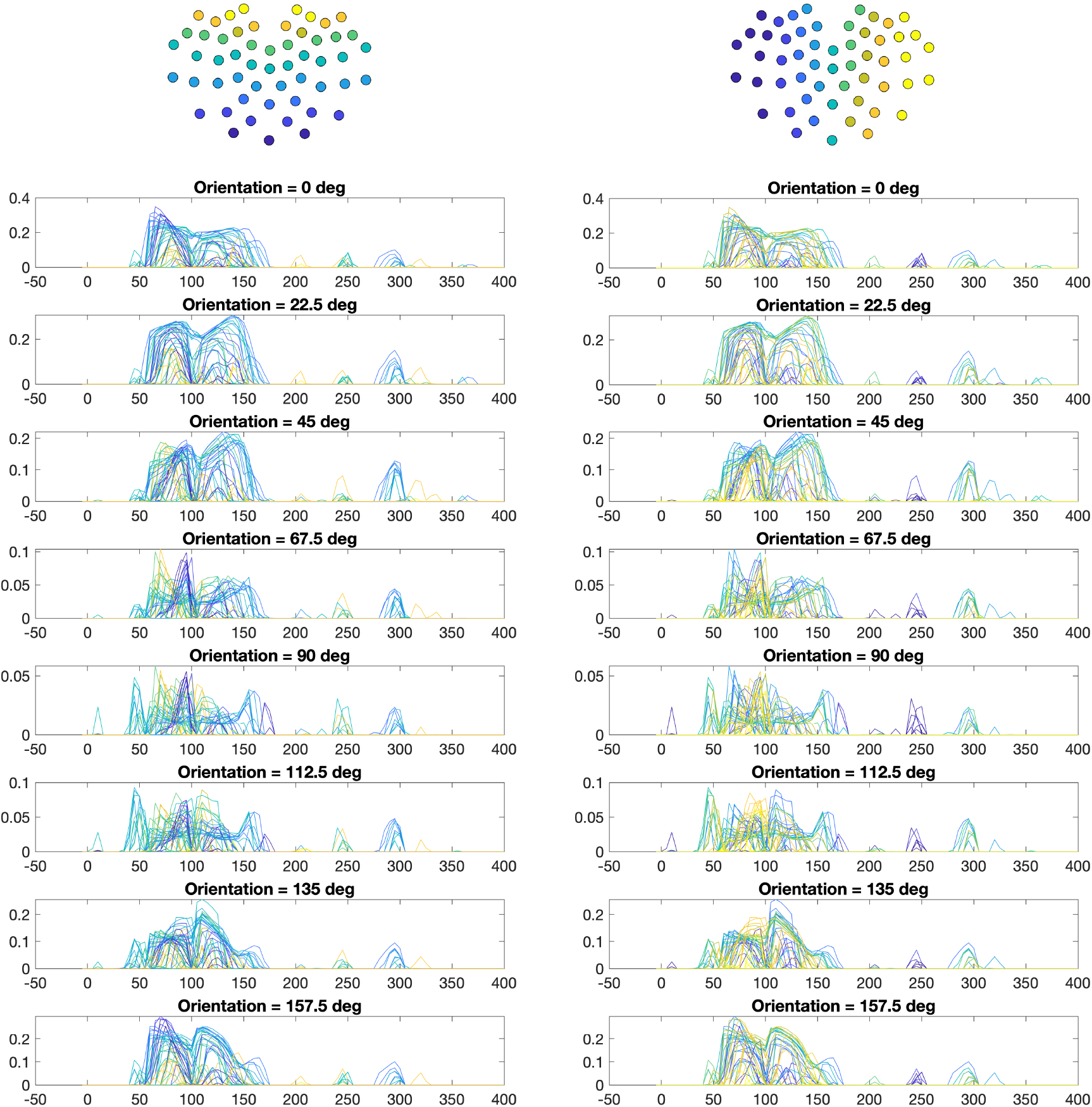
Results from the encoder tag probability over time analysis for the orientation tuned encoders. As with the SF probability over time analysis, we calculated the probability of observing pixels tagged with any given encoder’s peak orientation by summing the number of pixels tagged by each encoder for each electrode at each time point and then dividing each sum by the total number of visible pixels in the stimuli. Unlike the SF probability by time analysis, the orientation DETI mapping does not reveal any differences across the ventral-posterior to dorsal-posterior electrodes. Similar to the SF analysis, there is no clear lateralization of orientation probabilities. However, there is a tendency for the horizontally tuned encoders to be overall less prevalent than the other encoder orientations (note that the y-axes are different across encoder orientation). Please view the accompanying movie for a complete depiction of how different orientation DETI maps evolve over time [<u>**LINK**</u>].

**Supplementary Figure 4.**
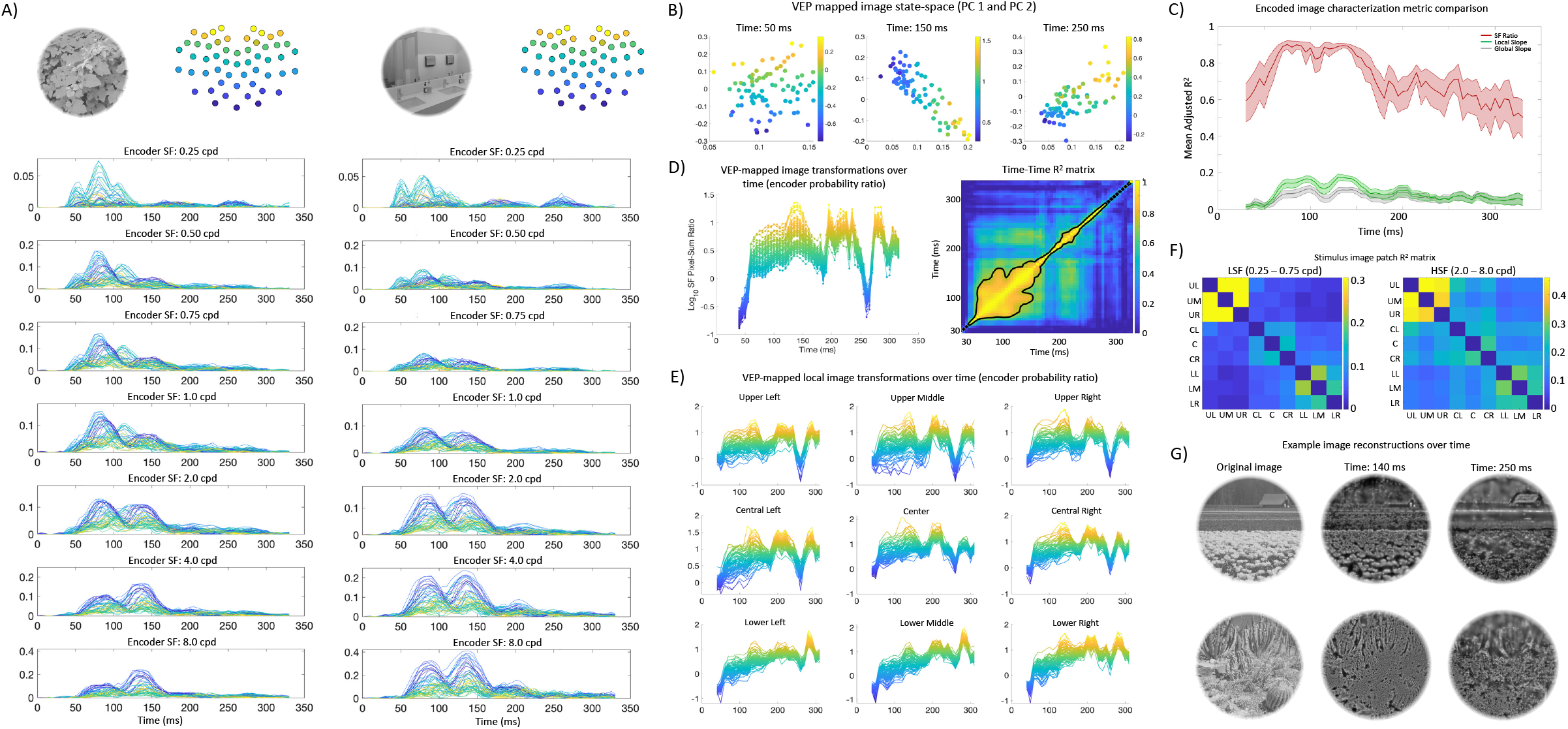
Image-specific DETI mapping procedure results using the Benjamini-Hochberg correction procedure with a false discovery rate of 5%. The layout of the results presented above is identical to that shown in **Figure 6** in the main article.

## Acknowledgements

James S. McDonnell Foundation grant (220020430) to BCH; National Science Foundation grant (1736394) to BCH and MRG.

## Notes

### Competing Interest Statement

The authors have declared no competing interest.

